# Capturing Spatially Organized Oscillatory Cliques as Signatures of Neuronal Assemblies

**DOI:** 10.64898/2026.03.14.711812

**Authors:** Katharina Heining, Pierre Le Merre, Mikael Lundqvist, Marie Carlén

## Abstract

State-of-the-art neural recording technologies now enable dense sampling of neuronal activity, demonstrating meaningful spiking variability from moment to moment. Yet spiking activity is sparse by nature and allows only partial access to collective neuronal dynamics. Local field potentials (LFPs) provide complementary information by capturing coordinated activity from large neuronal populations, but methods to characterize the fine-scale spatio-temporal organization of LFPs are lacking. Starting from the notion that neural oscillations are brief and burst-like, we introduce a framework to detect and analyze SPatially Organized Oscillatory Cliques (SPOOCs), oscillatory events that are cohesive in space, time, and frequency. SPOOCs displayed diverse dynamics in space and frequency and were differentially modulated during stimulus processing. Linking SPOOCs to local spiking, we demonstrate that these events index rapid reconfigurations of neuronal assemblies. With SPOOChunter, we provide an open-source toolbox that enables systematic detection and analysis of transient, spatially organized population dynamics in high-density electrophysiological recordings.

## Introduction

New technologies allow us to sample neural activity at ever-increasing density, but spiking activity is sparse by nature. In contrast, LFPs reflect instantaneous neuronal population activity and thus provide information about transient collective computations. However, methods to characterize the spatiotemporal dynamics of densely sampled LFPs are currently lacking. Here, we introduce SPatially Organized Oscillatory Cliques (SPOOCs), propose a framework for their detection and analysis, and provide evidence that SPOOCs reveal instantaneous formations of and changes in local spiking assemblies.

Recent work has revealed the brief and intermittent nature of oscillatory dynamics, often manifested as short, high-power bursts (Feingold et al., 2015; Lundqvist et al., 2016; Sherman et al., 2016; Kucewicz et al., 2017; Tinkhauser et al., 2017; Torrecillos et al., 2018). This understanding has led to a shift in how we characterize oscillations, beyond a coarse classification according to frequency band, taking characteristics such as specific frequency, duration, amplitude, and waveform into account (Douchamps et al., 2024; Lopes-dos-Santos et al., 2025). Studying the diversity of oscillations provides valuable insights into their differential function and mechanistic origin (Fernandez-Ruiz et al., 2023; Szul et al., 2023; Lundqvist et al., 2024). However, relatively few studies consider their spatial evolution (Zich et al., 2023). With SPOOCs we operationalize the concept of the diverse and dynamic nature of oscillations in space.

Here we utilize the dense sampling offered by Neuropixels to capture the fine-scale localization of oscillatory events and their impact on local spiking. Specifically, we define and extract SPOOCs as bursts of elevated power cohesive in space, time, and frequency. We then characterize the dynamics of individual SPOOCs in space and frequency and show how they change with behavior. By relating SPOOCs to local spiking, we reveal discrete assembly dynamics not detectable from spiking activity alone. With SPOOChunter we provide a toolbox for detecting and analyzing SPOOCs and their relationship to spiking, thereby increasing the information retrievable from multi-electrode recordings.

## Results

We detected SPOOCs from the LFPs of a single vertical column of Neuropixels probes, disregarding the second vertical column. We thus avoided zig-zagging in horizontal space and introducing a second spatial dimension (Fig. 1a). The vertical distance of 20 μm between successive channels dictates the SPOOCs’ spatial resolution. SPOOC extraction involves several steps, at the core of which is a channel-wise (N = 192 channels) spectral burst detection (Fig. 1b).

**Figure 1:**
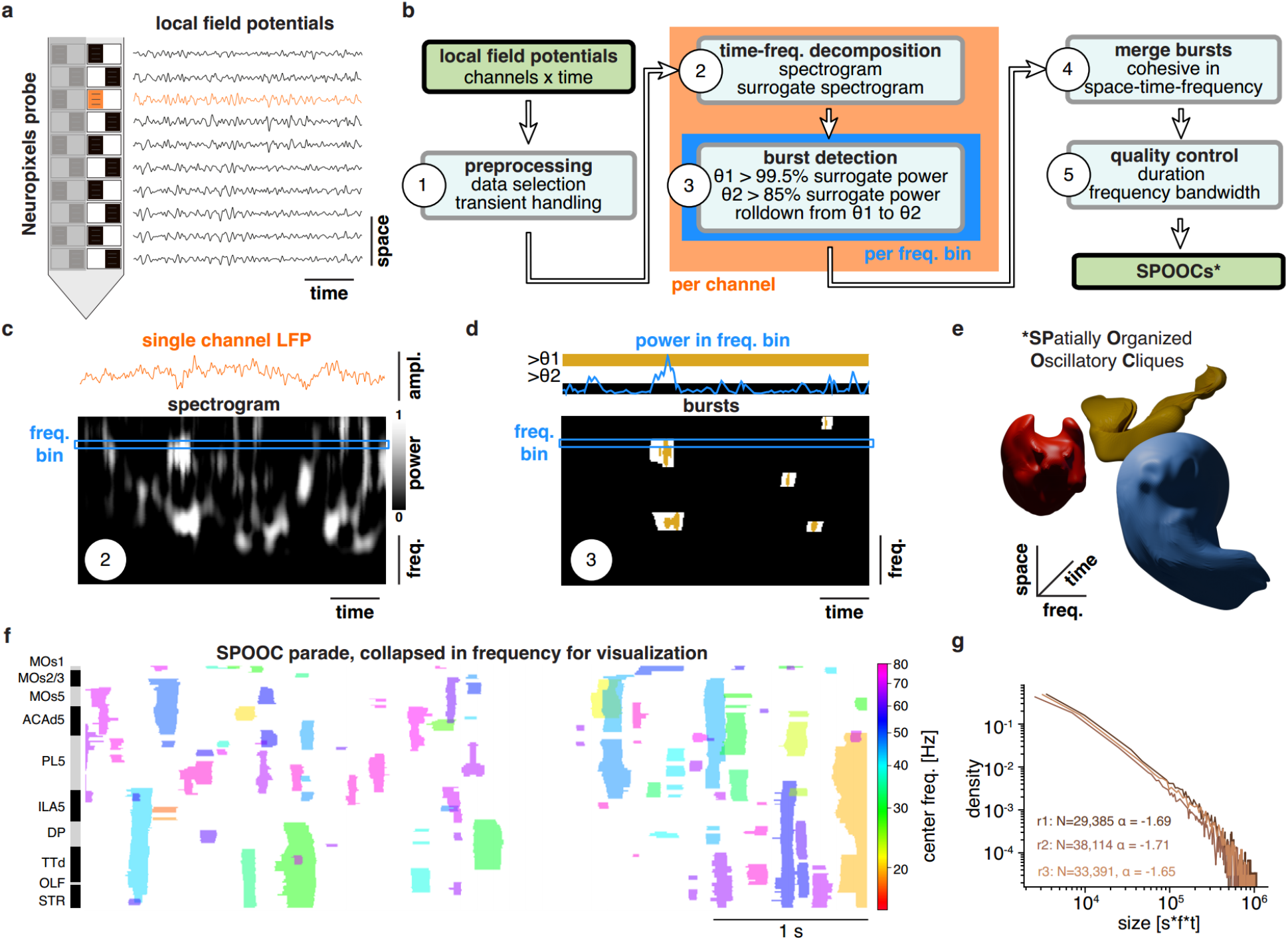
Detecting SPOOCs from multi-electrode LFPs. **a)** SPOOC detection operates on LFPs from one vertical column (N = 192 channels) of a Neuropixels probe. LFPs of eight example channels are shown; one channel is highlighted in orange. **b)** Flow diagram illustrating how SPOOCs are extracted from multi-channel probes. Highlighted are the channel-wise time-frequency decomposition (orange) and the burst detection per channel and frequency bin (blue). **c)** From each LFP (top, orange), spectrograms (bottom) are derived. To visually reveal power fluctuations per frequency bin, power is dynamically normalized per frequency bin (Waldert et al. 2013) to values between 0 and 1, corresponding to the 1^st^ and 99^th^ power percentiles, respectively. The blue box delimits an example frequency bin. **d)** *Top:* Power time series in an example frequency bin (blue) and thresholds θ1 (yellow) and θ2 (white) derived from the corresponding frequency bin of a surrogate spectrogram. Bursts are detected per channel and frequency as time intervals where the power continuously exceeds θ2 and at least once crosses θ1. *Bottom:* Burst spectrogram obtained by detecting bursts per frequency bin. Bursts are shown as white (power > θ2) and yellow (power > θ1) and present in this example as five burst entities cohesive in space and frequency. **e)** SPOOCs are obtained as power bursts cohesive in space, time, and frequency by stacking the burst spectrograms of subsequent channels. Colored 3D shapes schematically illustrate three SPOOCs. **f)** SPOOC sequence in space and time. For this 2D visualization, SPOOCs were collapsed in frequency and colored according to their center frequency. The spatial y-axis marks brain regions as alternating black and grey boxes. **g)** Size distribution of SPOOCs for the three recordings. SPOOC size was defined as the voxel count in the space-time-frequency volume. Linearity in the double logarithmic display indicates a power law. N: Total number of SPOOCs analyzed per recording; α: the exponent of the fitted power law.

### Preprocessing targets awake brain states and removes sharp transients

Preprocessing is essential to the SPOOC extraction procedure. To increase computational efficiency, we downsampled the LFPs to 500 Hz. We then detected and excluded episodes dominated by large, irregular amplitude activity. This type of activity occurs during sleep and drowsiness (Harris and Thiele, 2011) and would compromise identification of suitable power thresholds and hence spectral burst detection. In addition, we excluded episodes with salient artifacts due to stimulation or behavior, such as optogenetic stimulation or licking. A further problem are non-oscillatory transients, as they happen, for instance, during event-related potentials. In the spectral domain, such transients manifest as broadband high-power events. We detected and cut out these non-oscillatory transients for each channel individually and linearly interpolated between the data points bordering the gap. Preprocessing thus resulted in windows of active, awake activity that are cleared of sharp transients to facilitate the detection of genuine oscillatory bursts.

### Detecting spectral bursts per channel and frequency bin

After preprocessing, we bandpass filtered the LFPs using a Butterworth filter between 0.5 and 100 Hz, choosing 100 Hz as the upper limit for demonstration purposes and to limit spectral contamination by spiking activity. We calculated spectrograms for each LFP time series using a superlet transform (Fig. 1c). The superlet transform results from a geometric superposition of wavelets of multiple orders and thereby mitigates the resolution tradeoff between time and frequency inherent to spectral estimation (Moca et al., 2021). With the superlet transform, frequency bins can be arbitrarily small. For demonstration purposes and considering computational efficiency, we opted for a 1 Hz frequency resolution.

To identify thresholds suitable for detecting high-power bursts, we created surrogate LFPs using phase randomization, which preserves the overall power spectrum but destroys temporal phase structure (Feingold et al., 2015; Lancaster et al., 2018). From the spectrogram of a surrogate LFP, we identified two power thresholds per frequency bin: θ1 marking the 99.5th and θ2 marking the 85th percentile of the surrogate’s power distribution, respectively. The power time series of the corresponding channel and frequency bin in the original spectrogram was then thresholded using θ1 and θ2: Time intervals during which power values exceeded θ2 continuously and crossed θ1 at least once were defined and detected as bursts. Applying this burst detection procedure for each frequency bin resulted in channel-wise binarized burst spectrograms separating 2D bursts (frequency x time) from a non-burst background (Fig. 1d).

### SPOOCs are high-power bursts cohesive in space, time, and frequency

We obtained SPOOCs by stacking the channel-wise binarized burst spectrograms in space according to the original channel order and extracting cohesive 3D structures (Fig. 1e). Effectively, this amounts to aggregating all burst “voxels” in space-time-frequency, such that touching burst voxels constituted a SPOOC. A SPOOC can thus be understood as a high-power burst cohesive in space, time, and frequency. SPOOCs lasting less than 3 cycles of their average frequency (f_cent_) and SPOOCs with spectral bandwidths exceeding f_cent_ + 5 Hz were excluded. We further removed SPOOCs with f_cent_ below 15 Hz and above 80 Hz, respectively, in order to focus on oscillations in the beta and low-to mid-gamma range. SPOOC extraction resulted in a sequence of burst entities precisely defined in space, time, and frequency (Fig. 1f).

For further demonstrations, we selected three Neuropixels recordings with comparable probe placement mostly in the prefrontal cortex and good data quality (dataset DANDI:000473/0.230417.1502 from Calvigioni et al., 2023). Consistently across recordings, SPOOC sizes, defined by voxel count in the space-time-frequency volume, followed a slightly subcritical power law (Fig. 1g), as could be expected for subsampled systems close to criticality (Priesemann et al., 2009). The presence of a power-law size distribution indicates heavy-tailed dynamics characteristic of neuronal avalanches, originally observed for spiking activity (Beggs and Plenz, 2003; Chialvo, 2010).

### Categorizing and characterizing SPOOCs

To facilitate statistical analyses, we coarsely summarized SPOOCs into categories according to four features capturing their spectral extent and spatial positioning: each SPOOC’s upper and lower frequency bound and its top and bottom channel (Fig. 2a). Based on these four features, we clustered SPOOCs into nine categories, using the k-means algorithm. For our example recordings, this clustering led to an approximate tiling of the space-frequency plane into a 3-by-3 grid (Fig. 2b). SPOOCs of the same category displayed a considerable degree of individuality in terms of exact shapes (Fig. 2c). Accordingly, most analyses, e.g., relating SPOOCs to single-unit firing patterns, were carried out using the exact shape of individual SPOOCs. Generally, specific research questions will benefit from nuanced clustering strategies, possibly including duration, relative power, or 3D SPOOC shape. In this study, we aimed for as few and as simple categories as possible. In the following, we show that these suffice for demonstrating the general relevance of the SPOOC framework.

**Figure 2:**
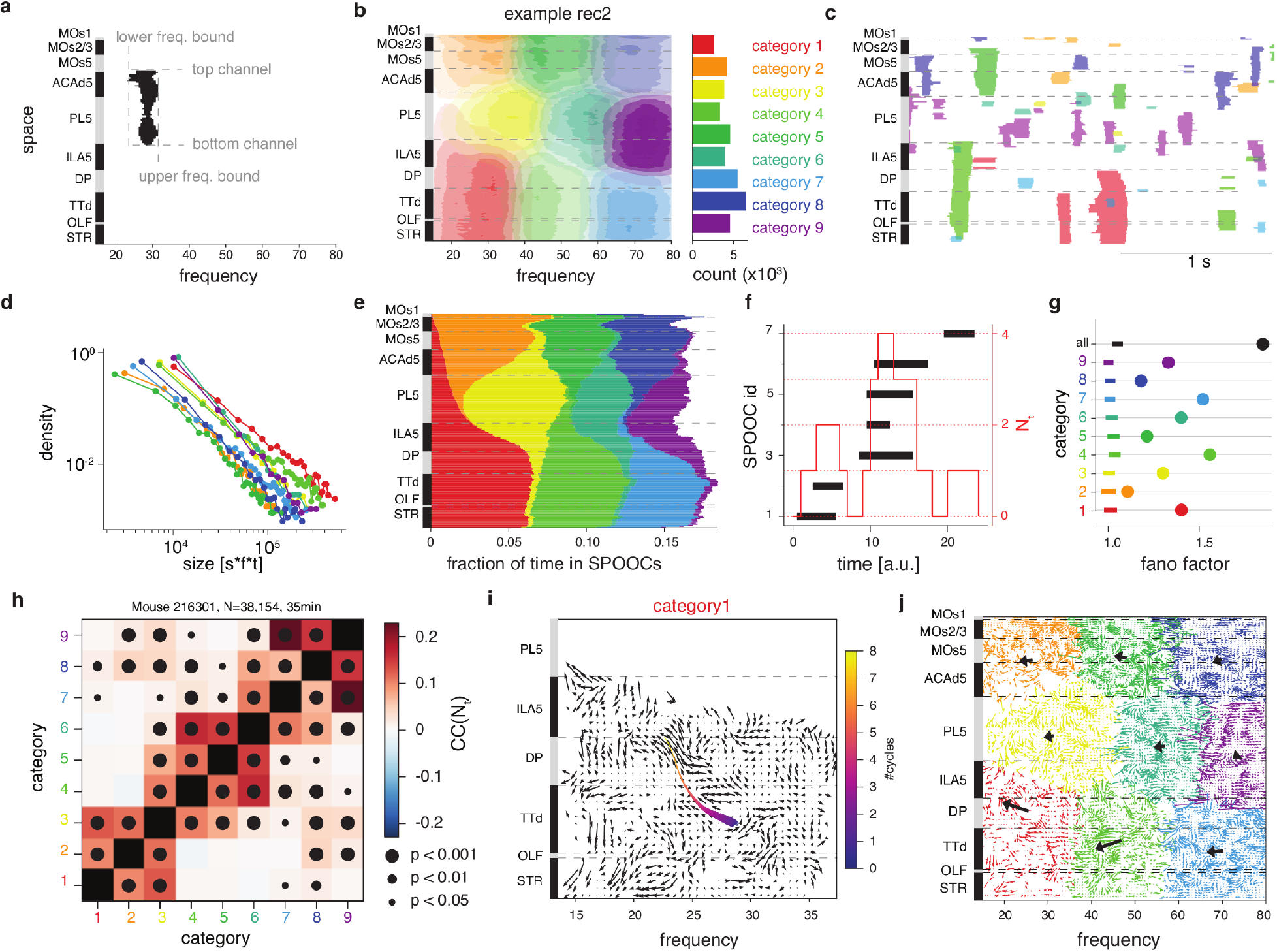
SPOOC characterization and dynamics. **a)** For clustering, each SPOOC was represented by four features: their lower and upper frequency bounds and their bottom and top channels. The black shape displays an example SPOOC collapsed in time. The spatial axis (left) marks brain regions as alternating black and grey boxes. **b)** *Left:* The SPOOCs of each category (colored) of an example recording are displayed collapsed in time, with the transparency of the shading indicating relative density levels. *Right:* Number of SPOOCs in each category. **c)** SPOOC sequence in space and time. For this 2D visualization, SPOOCs are collapsed in frequency and colored according to their category, as in b. **d)** Size distribution of SPOOCs per SPOOC category. SPOOC size was defined as the voxel count in the space-time-frequency volume. Linearity in the double logarithmic display indicates a power law. **e)** Fraction of time spent in SPOOCs per channel and SPOOC category displayed as a stacked bar graph. **f)** Schematic illustration of N_t_ (red), defined as the number of SPOOCs co-occurring at each point in time. The temporal extent of seven example SPOOCs is indicated by black horizontal bars. **g)** Temporal arrangement of SPOOCs. The Fano factor of N_t_ quantifies the temporal dispersion of SPOOCs; values above 1 indicate swarm-like clustering. Dots show Fano factors per SPOOC category (colors) and the overall Fano factor obtained by pooling SPOOCs across categories (black). Horizontal bars denote 90% confidence intervals derived from randomly distributing SPOOCs within analysis windows (maximum window duration: 40 s). **h)** Temporal co-occurrence of SPOOC categories. The Pearson correlation coefficient (CC) compares N_t_ (see g) between pairs of SPOOC categories. Dot size indicates significance level as indicated. High CC suggests that SPOOCs of two categories co-occur in time. **i)** Average spectrospatial flow field for SPOOCs of category 1. Black arrows indicate the direction and strength of propagation at each point in the space–frequency plane. The colored line (purple to yellow) shows the mean spectrospatial trajectory across SPOOCs, computed by averaging positions per oscillation cycle to accommodate variable cycle durations. Line width reflects the number of observations contributing to the trajectory and therefore decreases with cycle number. **j)** Average spectrospatial flow fields for all SPOOC categories (colors; see i). Black arrows at the spectrospatial center of each category indicate the grand-average propagation direction and magnitude.*Data:* All data shown in this figure originate from recording 2.

Overall, each channel spent around 16% of the recording time in SPOOCs, with variations across space (Fig. 2d). Spatial dips in SPOOC occurrence might be due to (1) natural spatial boundaries for rhythmically co-active populations, (2) layer boundaries where oscillations cancel to zero due to phase inversion, or (3) channels with low signal-to-noise ratio. We tried to mitigate the latter through spatial smoothing when merging spatially adjacent time-frequency bursts to SPOOCs. The size distribution of each individual SPOOC category followed a power law with varying exponents depending on SPOOC category (Fig. 2e). This suggests that the overall avalanche dynamics and critical behavior we observed (Fig. 1g) are replicated in each SPOOC category.

### SPOOCs come in spatial swarms of similar frequency

To analyze changes in SPOOC occurrence over time and motifs in the SPOOC sequence, we calculated N(t), the number of SPOOCs occurring simultaneously at any point in time (Fig. 2f). For all SPOOC categories, the Fano factors of N(t) ranged between 1.2 and 1.6 and significantly exceeded Fano factors expected from a random distribution of SPOOCs in time (Fig. 2g). This shows that SPOOCs of the same category occur in swarms or ‘superbursts’. Pooling SPOOCs from all categories results in an even larger Fano factor close to 2, demonstrating that SPOOC swarms consist of spectrospatially diverse SPOOCs and suggesting that the emergence of SPOOCs of diverse categories is governed by a common generative mechanism.

Asking whether certain SPOOC categories co-occur more often in time than others, we correlated N(t) between category pairs. To establish significance, we then compared the correlation coefficients to surrogates derived from versions of N(t) cyclically shifted in time. Generally, SPOOC categories of similar frequency that were distributed in space showed stronger and more significant correlations, while SPOOC categories with similar spatial location but different frequency characteristics displayed weaker and largely non-significant correlations (Fig. 2h). These findings indicate that there might exist a local competition between SPOOCs of different frequencies, preventing simultaneous occurrence. The clustering of SPOOCs of similar frequency across space might point to probe-wide (global) shifts in oscillatory regimes.

### SPOOCs slow down in frequency as they decay

Each individual SPOOC is defined by a particular 3D shape in the space-time-frequency volume. To analyze whether individual SPOOCs change their spectrospatial profile during their lifetime, we computed trajectories in the space-frequency plane for each SPOOC. These trajectories were obtained by temporally binning each SPOOC at the period duration of their central frequency and computing the mean in space and frequency during each temporal bin. The flow field created from the trajectories of all SPOOCs belonging to a certain category indicates for each point in the space-frequency plane the preferred direction of propagation (Fig. 2i). This allows, for example, to observe boundaries that SPOOCs do not tend to cross (unstable equilibria or sources) and to identify locations and frequencies SPOOCs tend to converge to (attractors in the space-frequency plane), thus providing valuable insights into the oscillatory dynamics *within* single SPOOCs. In our data, SPOOCs, especially those of the lower-frequency categories, tended to slow down in frequency over time (Fig. 2j). The flow field of higher-frequency SPOOC categories displayed finer substructures with vortex-like behavior, possibly suggesting the presence of dynamically distinct subcategories.

### SPOOCs significantly modulate single-unit firing

Oscillations, particularly in the gamma frequency range (30–100 Hz), are known to reflect changes in neuronal firing, both in terms of firing rate and spike timing (Fries et al., 2007; Vinck et al., 2013; Perrenoud et al., 2025). In being frequency-band agnostic and spatially specific, the SPOOC framework allows a nuanced characterization of the relationship between oscillatory activity and single-unit (putative single neuron) firing. First, we examined how firing rates are modulated during SPOOCs.

Does a single unit significantly change its firing rate during SPOOCs of a certain category? To evaluate this, we counted the number of spikes fired during time intervals in which SPOOCs of that category registered on the exact channel to which the unit was assigned (Fig. 3a). Only single units with ≥ 200 spikes overall and single-unit SPOOC pairings ≥ 20 s of a given SPOOC category available on the unit’s recording channel were analyzed. To establish significance, we compared the observed spike count to 100 surrogate spike counts obtained by cyclically shifting the spike train within temporal windows ≤ 40 s. A unit was considered significantly rate-modulated if the observed spike count during a given SPOOC category exceeded 95% (up-modulation) or fell below 5% (down-modulation) of the surrogate counts. Consistently across recordings and categories, SPOOCs modulated single-unit firing rates (Fig. 3b). Modulation probability varied strongly with SPOOC category. While all SPOOC categories displayed significant positive modulation probabilities, some SPOOC categories also exhibited strong down modulation of single-unit firing.

**Figure 3:**
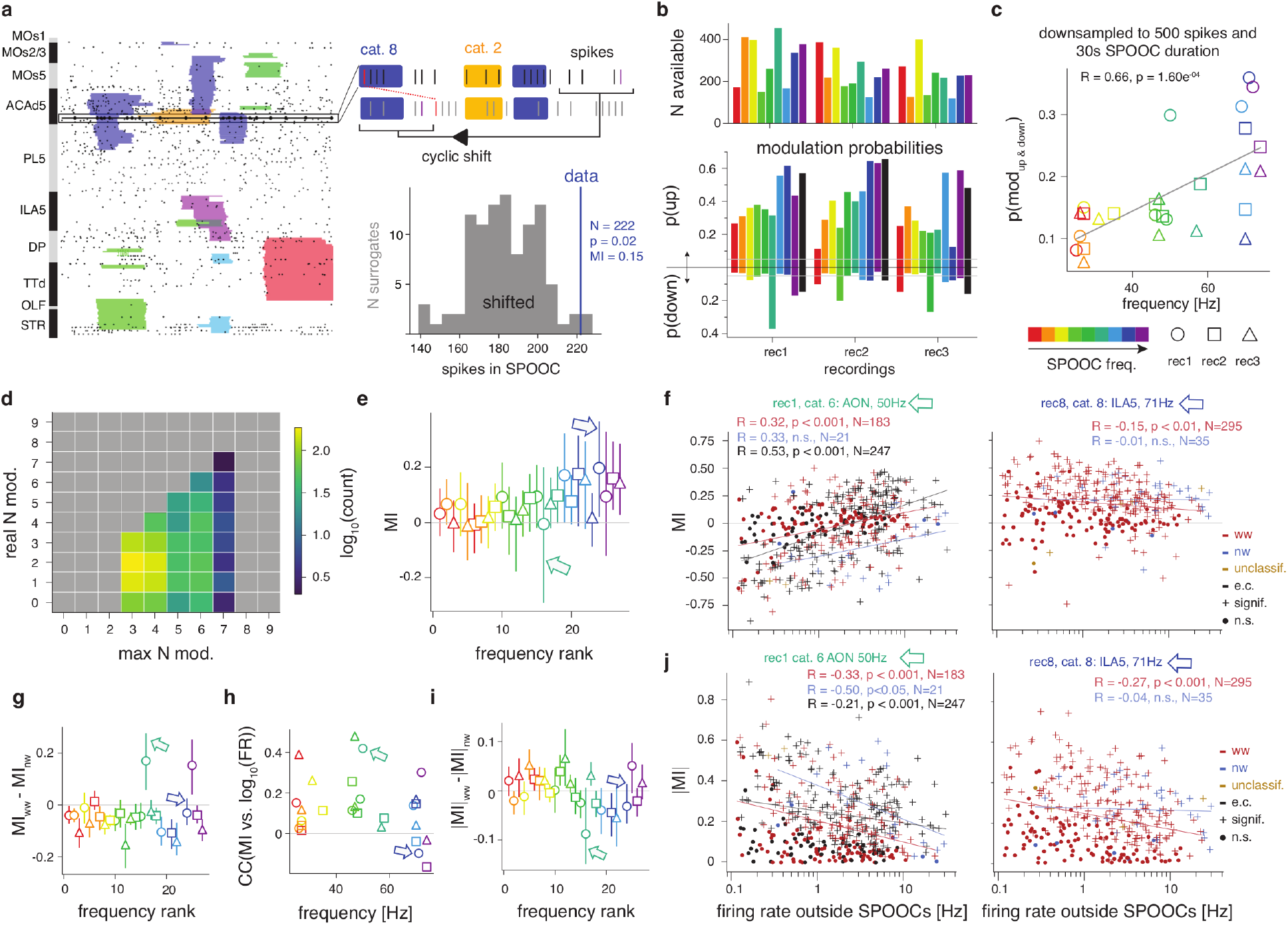
How SPOOCs modulate firing rates depends on SPOOC category and single-unit properties. **a)** *Left:* SPOOC sequence (colored shapes) and single-unit firing (black dots) across space and time. SPOOCs are collapsed across frequency and colored by category. The black box highlights the recording channel of a single unit, whose spikes are shown as large black dots connected by a line. *Top right:* Schematic of the per-channel evaluation of firing-rate modulation. During intervals in which a SPOOC of a given category (colored boxes) overlaps with the channel, spikes (black ticks) are counted. Surrogate spikes (grey ticks) are obtained by cyclically shifting the spike train by a random delay. Red ticks mark the first spike of the original sequence; dashed red line indicates the shift delay. *Bottom right:* Significance of rate modulation (p) is determined by comparing the observed spike count during SPOOCs of a given category (N, blue) with the distribution of counts from surrogate spikes (grey). The modulation index (MI) contrasts firing rates inside SPOOCs of a given category with the firing rate outside all SPOOCs. **b)** *Top:* Number of single units fulfilling the minimum data availability criterion (≥ 200 spikes and ≥ 20 s time in SPOOCs on the unit’s channel) per recording and SPOOC category. SPOOC categories are defined separately for each recording and ordered by average frequency from low (red) to high (purple). *Bottom:* Modulation probabilities, computed as the fraction of significantly up-modulated (positive values) and down-modulated (negative values) single units per recording and SPOOC category (colored bars). Black bars: overall locking probability to any SPOOC category per recording. Grey horizontal lines: 0.05 significance threshold. **c)** Combined probability of positive and negative firing-rate modulation as a function of average SPOOC frequency across categories. Colors indicate the frequency rank of each SPOOC category within a recording, and symbols denote recordings. For unbiased statistical evaluation, data were downsampled to 300 spikes and 30 s of SPOOC occupancy per single-unit SPOOC category pair. Statistics: Pearson correlation. **d)** Modulation of single units by SPOOCs of multiple categories. Two-dimensional histogram comparing the estimated maximum number of SPOOC categories significantly rate-modulating units (x-axis) with the observed number of significant rate modulators (y-axis). **e)** Distribution of modulation index per SPOOC category. Markers indicate the median modulation index across units, and vertical lines denote the 20th–80th percentile range. SPOOC categories are arranged on the x-axis by rank of average frequency. Colors and symbols as in c. Arrows indicate the SPOOC categories examined in detail in f. **f)** Modulation index as a function of firing rate outside SPOOCs (‘baseline firing rate’) across single units. The relationship is shown for two example SPOOC categories (left and right panel) and separated by unit type: wide-width (ww, red), narrow-width (nw, blue), and extra-cortical (ec, black). Regression lines and Pearson statistics are shown per unit type. Marker symbols indicate significance of modulation (dot: non-significant; plus: significant). **g)** Difference in mean modulation index between ww and nw units per SPOOC category. Markers indicate the mean; vertical lines show the 90% confidence interval obtained by bootstrapping (resampling units with replacement). SPOOC categories are arranged on the x-axis by rank of average frequency. **h)** Correlation coefficients between the decadic logarithm of baseline firing rate and the modulation index across single units per SPOOC category. SPOOC categories are arranged on the x-axis by average frequency. **i)** Difference in mean absolute modulation index between ww and nw units per SPOOC category. See g for details. **j)** Absolute modulation index as a function of firing rate outside SPOOCs across single units. See f for details.

We next assessed whether modulation probability correlated with SPOOC frequency. To avoid unequal amounts of data biasing significance statistics, we constrained the number of considered spikes to 300 and the time spent in SPOOCs of a given category to 30 s by randomly downsampling the data accordingly. Indeed, we observed a strong and highly significant positive correlation between modulation probabilities and average frequency of SPOOC categories (Fig. 3c), demonstrating that higher frequency SPOOCs have a higher probability of modulating single-unit firing rates.

Considering that modulation probabilities often exceeded 0.5 for individual SPOOC categories (Fig. 3b) and that most units spatially coincided with at least three different SPOOC categories (Fig. 2b), we expected single units to be significantly modulated by multiple SPOOCs. Indeed, comparing the maximum number of potentially modulating SPOOCs (per single unit) to the actual number of significantly modulating SPOOCs confirmed that single units are typically significantly modulated by one to three SPOOCs and that some single units reach the estimated maximum number of modulators (Fig. 3d). Note that this potentially underestimates the number of modulating SPOOCs due to our arbitrary data limit (≥ 200 spikes, ≥ 20 s time in SPOOCs, see above), which, given our surrogate-based significance statistics, might be insufficient to reveal small scale effects.

### Firing rate modulation depends on neuron and SPOOC characteristics

Having found that the firing rate of most single units is significantly modulated during SPOOCs (Fig. 3d), we next analyzed how the strength and directionality of this modulation varied with unit type and SPOOC category. For this, we defined a modulation index as MI = (νSPOOC − νfree) / (νSPOOC + νfree). The modulation index contrasts the firing rates inside a given SPOOC category (νSPOOC) with the firing rate outside of all SPOOCs (νfree). A modulation index close to 1 implies that νSPOOC >> νfree, a modulation index near zero that νSPOOC ≈ νfree, and a modulation index close to −1 that νfree >> νSPOOC. Thus, the modulation index expresses a combination of modulation magnitude and direction. νSPOOC was computed by counting all the spikes of a unit that occurred during SPOOCs of a category and dividing the count by the total time spent in that SPOOC category. νfree was obtained accordingly for SPOOC-free intervals. The median modulation index across units increased with SPOOC frequency (R = 0.73, p = 1.67×10^-5^), indicating that, on average, higher-frequency SPOOCs modulate single-unit firing more positively than low-frequency SPOOCs (Fig. 3e). However, we observed a large variation in modulation index across single-units for most SPOOC categories. To explain this diversity, we analyzed the modulation index separately for three different unit types: cortical wide-width units (ww), cortical narrow-width units (nw), and extra-cortical units (ec). Ww and nw units can be identified based on their spike waveform characteristics and have been associated with putative pyramidal neurons and inhibitory interneurons, respectively. In addition, we investigated the relationship between modulation index and the decadic logarithm of firing rate outside SPOOCs (lgFR). Indeed, both lgFR and unit type accounted for modulation index variability, and the nature of these unit-SPOOC interactions depended on SPOOC category (Fig. 3f). In general, across categories, SPOOCs tended to modulate nw unit firing rate more positively than ww unit firing rate (Fig. 3g). Moreover, we observed a consistent positive correlation between modulation index and lgFR, indicating that high-rate units get modulated more positively by SPOOCs than low-rate units (Fig. 3h). Notably, this correlation between modulation index and lgFR varied strongly across SPOOC categories, underscoring SPOOC heterogeneity.

We fitted a linear mixed model (LMM) to explicitly quantify the mutual interactions between modulation index, lgFR, and unit type while accounting for the variability across SPOOC categories (Table 1). The LMM confirmed that at average lgFR, nw units showed significantly stronger positive modulation than ww units and identified ec units to be less positively modulated than ww units. The effect of firing rate on ww and nw unit modulation was positive but not statistically significant, while firing rate had a strongly positive and significant effect on the modulation of ec units (compare Fig. 3f, left). Random-effect estimates indicated substantial variability across SPOOC categories both in terms of overall modulation at average firing rate and firing-rate dependence.

To specifically study modulation magnitude, we next analyzed the *absolute* modulation index. Low-frequency SPOOCs modulated ww units more strongly than nw units, while high-frequency SPOOCs mostly modulated nw units more strongly than ww units (Fig. 3i). This suggests that distinct SPOOC categories mark specific shifts in excitation-inhibition balance. Furthermore, the absolute modulation index was anti-correlated to firing rate (Fig. 3j), demonstrating a reduced gain or saturation effect in high-rate units. Together, these results highlight the functional diversity across SPOOC categories and indicate that firing-rate modulation during SPOOCs is jointly dependent on unit type and firing rate.

### Neurons lock to SPOOCs

How spikes are synchronized to oscillations has been extensively studied. For instance, during gamma oscillations, spikes are known to preferentially “lock” to the trough of the oscillation (Whittington et al., 2000; Fries et al., 2007; Buzsáki et al., 2012; Vinck et al., 2013; Kim et al., 2016). We investigated phase locking to SPOOCs in order (1) to validate SPOOCs as local signatures of synchronized, “locked” firing and (2) to explore whether and how locking differs across SPOOC categories and unit types.

We evaluated phase locking of single units to SPOOCs precisely at the channel a single unit was assigned to (Fig. 4a, left). Specifically, we (1) identified for each SPOOC its overlap interval with the channel of a single unit, (2) filtered the LFP in the range of the SPOOC frequency at that particular channel, (3) computed the phase of the filtered LFP in the overlap interval using the Hilbert transform, and (4) extracted the phases at the time points of single-unit spiking. Repeating this procedure for all SPOOCs of a given category, we collected all spike phases for each single-unit SPOOC category pairing (Fig. 4a, top right). We then measured phase-locking strength using pairwise phase consistency (PPC; Vinck et al. 2010). To establish significance of phase locking, we compared the original PPC to 100 surrogate PPC values obtained by randomly redistributing spikes within each overlap interval (Fig. 4a, bottom right). This ensured that the number of spike phases evaluated was the same in the original and surrogate data. A unit was considered significantly locked to a SPOOC category if its PPC exceeded 95% of surrogate PPCs. We also tested significance by deriving PPC values from cyclically shifted spikes (as illustrated in Figure 3a), yielding comparable results. Only single-unit SPOOC category pairings with at least 100 phase values, i.e., single units with at least 100 spikes occurring during SPOOCs of a given category, were considered for further analyses.

**Figure 4:**
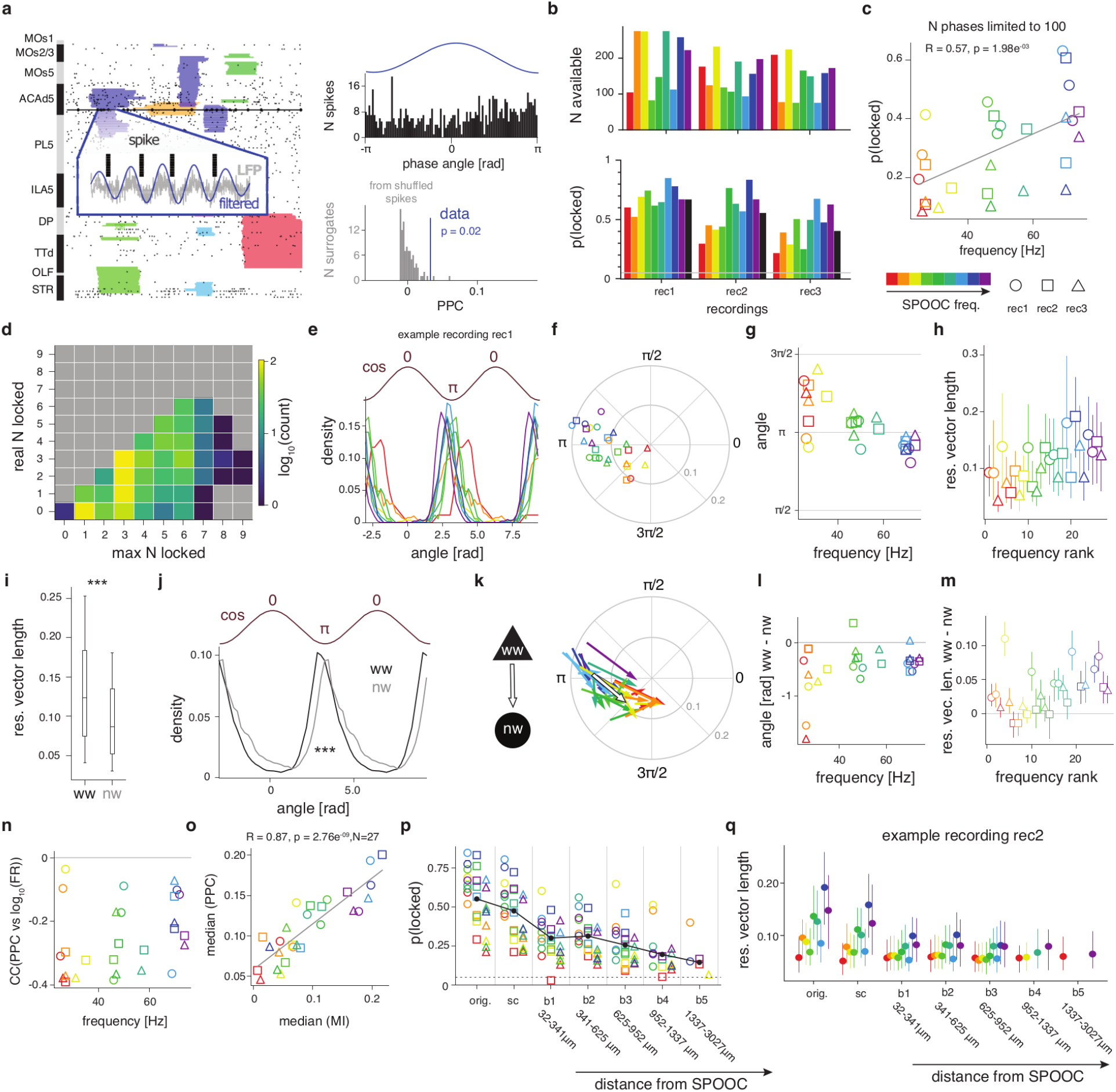
Phase locking to SPOOCs depends on SPOOC frequency, unit type, and distance. **a)** *Left:* SPOOC sequence (colored shapes) and single-unit firing (black dots) across space and time. Spikes of an example unit are shown as large black dots connected by a line. *Inset:* Time interval of a SPOOC overlapping the unit’s recording channel. The LFP recorded on the channel (grey) is filtered in the frequency range expressed by the SPOOC at that channel (blue). Phase values of the filtered LFP are extracted when the unit spikes (black ticks). *Top right:* Distribution of spike phases (black histogram) for an example unit during SPOOCs of a given category relative to the oscillation cycle (blue). *Bottom right:* Significance of locking (p) is determined by comparing the observed pairwise phase consistency (PPC, blue line) to surrogate PPC values (grey) obtained by randomly redistributing spikes within SPOOCs. **b)** *Top:* Number of single units fulfilling the minimum data availability criterion (≥100 phase values) per recording and SPOOC category. Categories are defined separately per recording and ordered by average frequency from low (red) to high (purple). *Bottom:* Locking probability, computed as the fraction of significantly locked units per recording and SPOOC category (colored bars). Black bars show the overall locking probability across SPOOC categories per recording. The grey horizontal line marks the 0.05 significance threshold. **c)** Locking probability as a function of average SPOOC frequency across categories. Colors indicate the frequency rank of each SPOOC category within a recording, and symbols denote recordings. For unbiased statistical evaluation, data were downsampled to 100 phase values per single-unit SPOOC category pair. Statistics: Pearson correlation. **d)** Single units lock to multiple SPOOC categories. Two-dimensional histogram comparing the estimated maximum number of SPOOC categories units could lock to significantly (x-axis) with the observed number of significant locking partners (y-axis). **e)** Density distribution of preferred phase angles across significantly locked units per SPOOC category (colors as in c) for an example recording. The brown curve above indicates the oscillation cycle for reference. **f)** Polar plot showing the average phase angle and locking strength (‘resultant vector’) per SPOOC category. Resultant vectors are obtained by representing each significantly locked unit as a unit-length polar vector and averaging these polar vectors across units. **g)** Average preferred phase angle as a function of average SPOOC frequency across categories. Circular correlation coefficient: 0.71. **h)** Distribution of locking strength (resultant vector length) per SPOOC category. Markers indicate the median across units, and vertical lines denote the 20th–80th percentile range. Categories are ordered by rank of average frequency. **i)** Comparison of locking strength between ww and nw units. The whisker plot displays the median (center line), interquartile range (25th–75th percentile), and the minimum and maximum non-outlier values (whiskers). Outliers, defined as values exceeding 1.5 times the interquartile range, are not shown. Statistics: Mann-Whitney U test, p = 1.32×10 ^-25^, N_ww_ = 2137, N_nw_ = 687. **j)** Density distribution of preferred phase angles for significantly locked ww (black) and nw (grey) units. The brown curve above indicates the oscillation cycle. Statistics: Watson-Williams test, p = 1.36×10^-27^. **k)** Polar plot contrasting resultant vectors between significantly locked ww and nw units per SPOOC category. Colored arrows connect the ww vector to the corresponding nw vector. **l)** Difference in mean preferred phase angle between ww and nw units per SPOOC category. Categories are sorted by average frequency. **m)** Difference in resultant vector length between ww and nw units per SPOOC category. Markers indicate the mean difference; vertical lines show 90% confidence intervals obtained by bootstrapping (resampling units per type with replacement). Categories are sorted by frequency rank along the x-axis. **n)** Correlation coefficient between the decadic logarithm of baseline firing rate and PPC across units per SPOOC category. Categories are sorted by average frequency. **o)** Correlation between median PPC and median rate-modulation index across SPOOC categories. Statistics: Pearson correlation. **p)** Distance dependence of phase locking per SPOOC category (colors, see c) and overall (black). Locking probabilities obtained from the precise overlap between SPOOCs and the unit’s channel (‘orig’, see a) are compared with probabilities computed using the full temporal and spectral extent of SPOOCs (‘full’, Methods), and for non-overlapping SPOOCs at increasing distances (bins b1–b4). Distances were adaptively defined and evaluated per single unit to ensure equal sample sizes (Methods). **q)** Distance dependence of locking strength (resultant vector length) for an example recording. Markers indicate the median across units, and vertical lines denote the 20th–80th percentile range. Binning (x-axis) as in p. SPOOC categories are arranged per bin by rank of average frequency.

The probability that a unit locked to a SPOOC of a certain category exceeded the expected 0.05 consistently and varied substantially across categories, with some SPOOC categories displaying phase-locking probabilities of > 0.7 (Fig. 4b). To avoid biases introduced by unequal amounts of data when evaluating significance statistics, we downsampled the phase values to 100 when assessing the relationship between SPOOC frequency and phase-locking probability. Phase locking probability showed a strong and significant positive correlation with SPOOC frequency (Fig. 4c), indicating that single units lock with higher probability to SPOOCs of higher frequency.

### Individual neurons can lock to multiple SPOOCs

The high probability of phase locking observed for individual SPOOC categories (Fig. 4b) suggested that single units could lock to more than one SPOOC category. Indeed, comparing the maximum number of SPOOC categories a unit could lock to with the actual number, units often achieved the maximum number of expected locking partners (Fig. 3d). For instance, the population of units that could lock to maximally three SPOOCs was split quite evenly between units locking to 1, 2, or 3 SPOOC categories. Note that these estimates are conservative, as we defined the maximum number of SPOOC categories a unit could lock to simply through data availability (at least 100 phase values), and small effects require more data to be detected as significant. Overall, the flexible adjustment of spike timing to multiple SPOOCs is in line with the local competition observed between SPOOCs of different frequency characteristics (Fig. 2h).

### Neurons fire earlier in the trough of high-frequency SPOOCs

We next examined at which phase of SPOOCs units preferred to fire. For this, we expressed each phase value, i.e., each oscillation phase at which a spike was fired, as a unit-length vector and computed the average phase vector across all unit vectors of each single-unit SPOOC category pair. The angle of this resultant vector gives the preferred phase, and the length indicates, similarly to the PPC, phase-locking strength. As expected, units preferred to fire during the trough of the SPOOC oscillations (Fig. 4e,f). Interestingly, however, we observed a significant anti-correlation between phase angle and SPOOC frequency (circular correlation coefficient Rcirc = 0.71; Fig. 4g), as well as a significant positive correlation between median resultant vector length and SPOOC frequency (R = 0.6, p = 1.05×10-3; Fig. 4h). Put simply, units locked stronger and fired earlier in the trough of higher-frequency SPOOCs. The observation that stronger locking is associated with earlier firing agrees with the idea that more activated units fire earlier in the oscillatory cycle (Fries et al., 2007).

### Phaselocking to SPOOCs depends on unit type and firing rate

Having established a SPOOC category-specific influence on single-unit spike timing, we next asked whether different unit types were differentially impacted by SPOOCs. Overall, we found that ww units locked significantly stronger (Fig. 4i) and fired significantly earlier in the phases of SPOOCs (Fig. 4j). Critically, this relationship was preserved across SPOOC categories (Fig. 4k): ww unit firing preceded nw unit firing consistently across categories (Fig. 4l), and the locking strength of ww units was consistently higher than the locking strength of nw units, with more pronounced unit-type differences during high-frequency SPOOCs (Fig. 4m). Our observation that ww units precede nw units is consistent with the proposed pyramidal-interneuron-gamma (PING) mechanism, where pyramidal neurons drive inhibitory interneurons, which in turn synchronously inhibit the pyramidal neurons, giving rise to the gamma rhythm (Whittington et al., 2000; Tiesinga and Sejnowski, 2009).

The correlation between outside-SPOOC firing rate (lgFR, see above) and PPC was strongly negative for all SPOOC categories (Fig. 4n), indicating that rate units with lower firing rates adjust their timing more precisely to SPOOCs. Moreover, the median PPC correlated strongly with the median modulation index across SPOOC categories (Fig. 4o), demonstrating that units lock more strongly (high median PPC) to SPOOCs that are more positive rate modulators (high median modulation index). This suggests that SPOOCs preferentially entrain units whose firing they modulate strongly, possibly ‘binding’ the excess spikes evoked.

### Phase locking is spatially confined to SPOOC boundaries

So far, we have introduced SPOOCs as entities defined in time, frequency, and space and shown that neurons lock to SPOOCs with high probability, suggesting that SPOOCs reflect synchronized neuronal firing patterns. To investigate whether the spatial extent of SPOOCs indicates spatially constrained firing of synchronized neuronal assemblies, we examined how phase locking changed with distance from the SPOOC location. To avoid biases arising from unequal amounts of data at different distances, we designed an adaptive spatial binning algorithm that ensured comparable sample sizes across bins (Methods).

We observed the highest locking probabilities for the original evaluation method, where phase locking was evaluated at the exact time intervals when SPOOCs overlapped with the single units’ recording channel (Fig. 4p). Overall locking probability slightly decreased when the full spectrotemporal extent of overlapping SPOOCs was used for evaluation (‘full’ condition; see Methods) and dropped markedly already in the first outside spatial bin. From the third spatial bin onwards (starting at 625 μm distance from SPOOCs), locking probabilities continuously dropped. The distance-dependent drop in locking probability was strongest for high-frequency SPOOC categories. Analyzing locking strength as a function of distance from SPOOCs further confirmed that phase locking is strongest locally for high-frequency SPOOCs (average frequency >50 Hz) and that consequently the most dramatic drop in locking strength occurs for high-frequency SPOOCs (Fig. 4q). For lower-frequency SPOOC categories, in contrast, locking strength hardly changed across distance. This suggests that high-frequency SPOOCs are signatures of locally synchronized firing, while low-frequency SPOOCs reflect more global rhythmic patterns.

### SPOOC rates differentially change depending on behavioral context

Thus far, we have established SPOOCs as entities with individual dynamics (Fig. 2) and as signatures of neuronal firing patterns (Figs. 3 and 4). A description of how SPOOCs change with time and depending on behavior would hence allow us to infer changes in coordinated single-unit firing. The recordings analyzed in the current study stem from a dataset using an aversive conditioning task (Calvigioni et al., 2023). In brief, experiments contained two blocks with auditory stimulation (Fig. 5a). During a passive listening block (100 trials), auditory stimulation consisted of a 200 ms 10 kHz tone stimulus. During a subsequent aversive block (50 trials), a 200 ms blue noise stimulus was followed by an aversive air puff to the eye. Individual trials were separated by inter-trial intervals (5–10 s; randomly picked from a uniform distribution).

**Figure 5:**
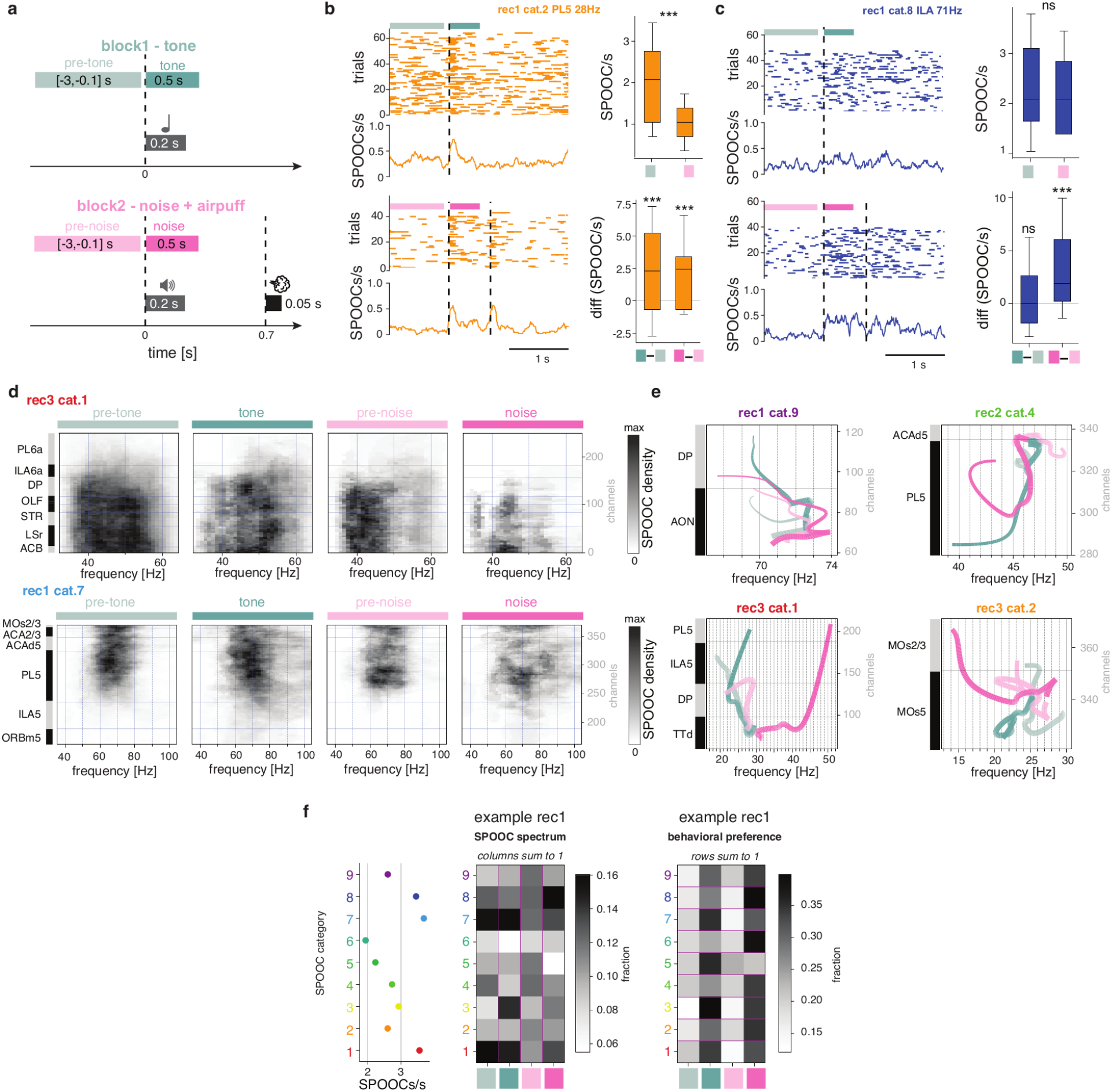
SPOOC rates and dynamics change depending on behavioral context. **a)** Structure of the behavioral paradigm. In the first block (teal, top), 200 ms pure tones were presented. In the second block (pink, bottom), 200 ms blue noise stimuli were followed by an aversive air puff. Four time periods were analyzed (colored horizontal bars): pre-tone, tone, pre-noise, and noise. Text on the colored bars indicates the respective analysis intervals relative to sound onset. **b)** Occurrence of an example SPOOC category across behavioral contexts. *Top left:* Interval raster plot displaying the timing and duration of individual SPOOCs (orange bars) across trials aligned to tone onset (vertical dashed line). The horizontal extent of each bar indicates SPOOC onset and offset. Teal bars mark the pre-tone (lighter shade) and tone analysis intervals. The orange curve below shows the trial-averaged occurrence rate. *Bottom left:* Same representation aligned to noise onset. Pink bars mark the pre-noise (lighter shade) and noise analysis intervals. *Top right:* SPOOC rates during the pre-tone and pre-noise intervals across trials. The whisker plot displays the median (center line), interquartile range (25th–75th percentile), and the minimum and maximum non-outlier values (whiskers). Outliers, defined as values exceeding 1.5 times the interquartile range, are not shown. Statistics: Mann-Whitney U test. *Bottom right:* Differences in SPOOC rate between tone and pre-tone windows (left whisker plot) and between noise and pre-noise windows (right whisker plot). Statistics: Wilcoxon signed-rank test. ***c)*** *Same as b, but for a different SPOOC category*. **d)** Spectrospatial profiles of two example SPOOC categories (top and bottom rows) for each behavioral context (columns). SPOOCs occurring in each behavioral context were collapsed across time and summed to produce a relative density profile (grayscale) in the space–frequency plane. To ensure comparability, the number of SPOOCs contributing to each panel was randomly downsampled to match the behavioral context with the fewest observations (top: N = 32; bottom: N = 78). **e)** Spectrospatial dynamics of four example SPOOC categories across behavioral contexts. Each colored line shows the mean spectrospatial trajectory across SPOOCs, computed by averaging spectrospatial positions per oscillation cycle to accommodate variable cycle durations. Line width reflects the number of observations contributing to the trajectory and therefore decreases with cycle number. **f)** Relative occurrence of SPOOC categories across behavioral contexts. *Left:* SPOOC rate per category across all behavioral windows combined (see a). *Center:* SPOOC spectrum showing the relative fractional count (grayscale) of each SPOOC category (rows) per behavioral context (columns). Each column sums to 1. *Right:* Behavioral context preference per SPOOC category. Each row shows the relative occurrence of a SPOOC category across behavioral contexts. Rows sum to 1.

To analyze changes in SPOOC occurrence during behavior, we represented each SPOOC as a time interval, generating one interval time series per SPOOC category. Peri-event time histograms revealed considerable diversity across SPOOC categories: for example, SPOOCs of certain categories strongly and transiently increased their rate following both types of auditory stimulation and the air puff, while generally decreasing their baseline rate during the aversive block (Fig. 5b). Other SPOOC categories showed no changes in baseline rate across blocks or in response to the tone stimulus, but a prominent sustained rate increase specifically following the onset of the noise stimulus announcing the aversive air puff (Fig. 5c). This diversity suggests that different SPOOC categories may index distinct network processes engaged during sensory processing and aversive anticipation.

### Behavioral context alters SPOOC substructure

The observed category-specific change in SPOOC occurrence during behavior, combined with our coarse categorization of SPOOCs, led to the hypothesis that SPOOCs of the same category exhibit more subtle systematic differentiations depending on behavioral context. We therefore investigated whether SPOOCs of the same category differed in shape depending on the behavioral context they occurred in. SPOOCs of certain categories strongly refined their spectrospatial profile during the aversive block and most prominently in the 500 ms following the onset of the noise stimulus announcing the air puff (Fig. 5d, top). In contrast, other SPOOC categories displayed a spectrospatial expansion in response to both types of auditory stimulation (Fig. 5d, bottom). This demonstrates that SPOOC categories are heterogeneous in how they change shape in relation to behavioral context and information processing.

Building on our previous finding that the spectrospatial signature of SPOOCs evolves during the lifetime of individual SPOOCs (Fig. 2i,j), we studied how the SPOOCs’ space-frequency trajectories differed across behavioral contexts (Fig. 5e): While some SPOOC categories displayed similar space-frequency trajectories across behavioral contexts, we noted a divergence of trajectories in other categories. Most prominently, trajectories during the noise announcing the air puff tended to diverge from the pre-stimulus trajectories.

### Behavior reshapes the SPOOC spectrum

When SPOOCs change their shape, i.e., their profile in space, time, and frequency, they may eventually transition between categories depending on the clustering resolution. We tracked systematic shifts in the relative abundance of SPOOC categories through a *SPOOC spectrum*, a space-frequency spectrum of oscillatory activity (Fig. 5f). The SPOOC spectrum shows how different SPOOC categories prevail at different times. In our data, these abundance patterns shifted across behavioral contexts: some SPOOC categories became relatively more prominent during auditory stimulation or during the aversive block, respectively, whereas others occurred more uniformly across behavioral contexts. Conversely, certain relatively rare SPOOC categories appeared selectively during specific behavioral contexts. Interpreting SPOOCs as signatures of locally coordinated neuronal firing, these shifts in the SPOOC spectrum point to changes in the participation of neuronal assemblies across behavioral contexts.

## Discussion

We have introduced SPOOCs as high-power bursts cohesive in space, time, and frequency. SPOOCs manifest in LFPs of high-density electrophysiological recordings and allow capturing a great diversity of intermittent spectrospatial dynamics. This framework unearthed previously hidden neural dynamics from high-density recordings and enabled refined single-trial assembly analyses.

Importantly, when extracting SPOOCs, we do not confine the analysis to predefined frequency bands but rather perform the extraction at high-frequency resolution. This results in SPOOCs with precise temporal and spectrospatial organization. Critically, to remain sensitive to precise organization, analyses were performed on individual SPOOC instances, e.g., when analyzing the relationship between SPOOCs and single-unit firing. SPOOCs were clustered into categories only to summarize and visualize results, revealing meaningful SPOOC-specific differences at this coarse level. We intuit that clustering tailored to more detailed features of SPOOCs will allow researchers to target more specific questions.

Transient bursts of oscillatory activity, particularly in the beta and gamma range, have previously been shown to influence the rate and variability of local spiking (Lundqvist et al., 2022; Boroujeni et al., 2023; Perrenoud et al., 2025; Harris et al., 2026). Consistent with these observations, SPOOCs impacted both the firing rate and spike timing of individual neurons. The impact depended both on SPOOC category and neuron type. Overall, nw neurons were more positively rate-modulated by SPOOCs than ww neurons, whereas ww neurons showed stronger phase-locking than nw neurons. In addition, ww neurons fired earlier in the SPOOC cycle, consistent with excitation-inhibition-generated oscillations (Whittington et al., 2000; Tiesinga and Sejnowski, 2009). High-frequency SPOOCs boosted and phase-entrained neuronal firing more strongly and locally than low-frequency SPOOCs. Phase-locking was strongly distance-dependent, establishing SPOOCs as spatially precise signatures of assembly activation. The overall heterogeneity of SPOOC–neuron relationships and the fact that individual neurons were influenced by multiple SPOOC categories suggest the existence of multiple overlapping assemblies with distinct properties.

Further, observing assembly activations through SPOOCs opens new insights into the nature of assemblies. Traditionally, assembly detection relies on identifying precise repetitions of spike time patterns and is, for statistical reasons, biased towards high-rate neurons, which we found to be least modulated by SPOOCs. In contrast to spike-based assembly detectors, assembly observation through SPOOCs does not demand the neurons to repeat their exact spiking or to even be active during each assembly occurrence. Leveraging LFPs provides a dense sampling of coordinated activity where the overall characteristics of the assemblies, their exact timing, and spatial extent are revealed.

Since SPOOCs reveal assembly activations, they allow us to infer large-scale assembly dynamics. For instance, SPOOCs display power law size distributions and swarming behavior reminiscent of the critical dynamics reported for spiking activity (Beggs and Plenz, 2003; Priesemann et al., 2009). This suggests that assemblies, defined as coherently oscillating groups of neurons, also display these behaviors. In addition, SPOOCs tended to slow down in frequency during their lifetime, suggesting that also the activity of the associated assemblies gradually slowed down toward the end of SPOOC events. Such progressive slowing is consistent with critical slowing down, a phenomenon observed in dynamical systems approaching a transition (Scheffer et al., 2009). We have further demonstrated that SPOOCs can be related to behavior. In principle, SPOOCs can be related to any functional parameter of choice, such as time-varying pupil size, whisking, or breathing.

In sum, we have shown that LFPs from high-density electrophysiological recordings can be used to infer neural assembly dynamics and abrupt changes in population activity, both in terms of large-scale dynamical properties and single neuron participation. Going forward, studies linking SPOOCs with cognitive processes are therefore particularly interesting. In addition, optogenetic experiments paired with multi-region recordings (as suggested in Fernandez-Ruiz et al., 2023) would shed light on the mechanisms of SPOOC generation and thereby assembly formation. Thus, SPOOC analysis, as a link between single neuron and population activity, provides insight into the precise spatiotemporal evolution of neural computations.

## Methods

### Evaluation of distance-dependent phase locking using spatially adaptive binning

First, we computed the instantaneous phase of each SPOOC. For this, we filtered the LFP recorded at the spatial center of the SPOOC within the SPOOC’s frequency range and over its full temporal extent and obtained the instantaneous phase using the Hilbert transform. Each individual SPOOC *i* was thus represented as a phase time series φ_i_(t), defined between the start and end of the SPOOC.

Next, we quantified the spatial relationship between units and SPOOCs. For each recording channel, we calculated the minimal distance to the spatial boundary of the SPOOC (e.g., the distance to the nearest SPOOC channel). This channel–SPOOC distance defined the distance between all units recorded on that channel and the corresponding SPOOC.

We then extracted spike phases for all single-unit SPOOC pairs in which the unit did not spatially overlap with the SPOOC by sampling φ_i_(t) at the spike times of the unit.

To avoid biases arising from unequal data amounts at different distances, we used an adaptive spatial binning procedure that ensured comparable sample sizes across bins. For each single-unit SPOOC category pair, we used the number of spike phases available during the exact spatial overlap between the unit’s channel and the SPOOC as a reference sample (see Fig. 4a). We then constructed successive distance bins such that each bin contained the same number of phase values as this reference sample. Because SPOOC occupancy and spike timing differed across channels and units, the resulting bin widths varied across single-unit SPOOC pairs.

To aggregate results across units, we defined common spatial bins that contained data from at least 50 single units per SPOOC category. When multiple unit-specific bins overlapped a given common bin, we selected the bin whose center was closest to the center of the common bin.

Finally, to test whether the precise spatial shape of SPOOCs influenced phase locking, we collected phase values directly from φ_i_(t) for spatially overlapping single-unit SPOOC pairs (‘full’ condition).

## Data and code availability

The toolbox SPOOChunter, comprising preprocessing, SPOOC extraction, and analysis of single-unit–SPOOC interaction, is available at [github-link: inserted on publication]. Documentation is provided at [inserted on publication]. The code used to generate all manuscript figures is available at [zenodo-link: insert on publication]. The data from the three example Neuropixels recordings is available for download in Neurodata Without Borders (NWB) format on DANDI Archive: https://dandiarchive.org/dandiset/000473 and https://dandiarchive.org/dandiset/001260.

## Acknowledgments

We thank Arvind Kumar and Ida Välikangas Rautio for insightful discussions and Jacob Langeloh for semantic and conceptual feedback. This work was supported by Wallenberg Scholar program (Knut and Alice Wallenberg Foundation), a KAW project grant (Knut and Alice Wallenberg Foundation), the WennerGren foundation, Hjärnfonden, ERC starting grant 949131, VR 2022-02328, a NARSAD Young Investigator Grant (Brain & Behavior Research Foundation) and a Stratneuro Postdoctoral Grant.

